# Privatisation rescues function following loss of cooperation

**DOI:** 10.1101/326165

**Authors:** Sandra B. Andersen, Melanie Ghoul, Rasmus L. Marvig, Zhuo-Bin Lee, Søren Molin, Helle Krogh Johansen, Ashleigh S. Griffin

## Abstract

A single cheating mutant can lead to the invasion and eventual eradication of cooperation from a population. Consequently, cheat invasion is often considered as “game over” in empirical and theoretical studies of cooperator-cheat dynamics, especially when cooperation is necessary for fulfilling an essential function. But is cheat invasion necessarily “game over” in nature? By following a population of bacteria through loss of cooperation and beyond, we observed that individuals evolved to replace cooperation with a selfish, or “private” behaviour. Specifically, we show that when cheating caused the loss of cooperative iron acquisition in a collection of *Pseudomonas aeruginosa* isolates from cystic fibrosis patients, a private uptake system that only benefits the focal individual was upregulated. This observation highlights the importance of social dynamics of natural populations and emphasizes the potential impact of past social interaction on the evolution of private traits.

## Introduction

Identifying mechanisms responsible for the maintenance of cooperation is a major achievement of evolutionary biology (Axelrod & Hamilton, 1981; Hamilton, 1963). We can predict in which conditions cooperation will thrive, and where it might pay to exploit cooperative neighbours. Evidence of tension between cooperative and cheating strategies are all around us in nature-for example, the co-evolution between flowers and their pollinators, the policing and counter-policing behaviours of social insects (Foster & Ratnieks, 2000) - and in human society (Mathew & Boyd, 2011; Rand et al., 2011). Cases where these dynamics have resulted in the loss of cooperation are much less well understood, primarily because long-term consequences of cheat invasions are unobserved and unreported (Sachs & Simms, 2006). This is for the obvious reason that it is inherently difficult to detect a history of something that no longer exists. And also because it is generally the case that once cheating has gone to fixation, it is almost impossible for cooperation to re-invade (Axelrod & Hamilton, 1981). The cheats won. End of story.

Life, however, goes on. Consider a population where cheating has gone to fixation. If cooperation fulfilled an important function, this population may now be at risk of extinction (Fiegna & Velicer, 2003). To escape this fate, selection may favour individuals that can restore function by employing a “privatisation” strategy - replacement of a mechanism that was once performed cooperatively as a group, with a selfish one that only benefits the actor (Bel, 2006). As such, we use the term privatisation as it is normally understood in common language to describe a switch in strategy from one that helps a whole group to achieve a goal to one where a goal is achieved by the actor alone. This differs from its use for describing the acquisition of property for future benefit by a group or an individual (Strassmann & Queller, 2014), the retention of public goods for individual use (Asfahl & Schuster, 2017), and for when public goods only are useable to a part of a community (Niehus et al., 2017). Privatisation could be a common occurrence in nature that has nevertheless been overlooked because there is no reason, *a priori*, to interpret private function as a result of past social interaction. It may also have been missed for the practical reason that we are required to track behaviour over many generations post-cheat invasion in a natural population. We overcome these difficulties by studying the evolutionary dynamics of a cooperative trait for more than an estimated 20 million generations (Table S1), in a population of bacterial cells. We report the first observation (to our knowledge) of a natural population responding to cheat invasion by adopting a privatisation strategy and, therefore, avoiding the possibility of extinction.

Our study population is comprised of *Pseudomonas aeruginosa* bacteria causing lung infection in patients with cystic fibrosis (CF). CF is a genetic disease that causes the build-up of dehydrated mucus in lungs and sinuses, which *P. aeruginosa* infects (Folkesson et al., 2012). The host actively withholds iron to limit infection, and, as a counter-measure, *P. aeruginosa* employ a range of different iron acquisition strategies (Poole & McKay, 2003). The primary mechanism relies on secretion of the siderophore pyoverdine (Andersen et al., 2015; Haas et al., 1991; Konings et al., 2013). This has been demonstrated to be a cooperative trait, where iron-bound pyoverdine molecules are available for uptake not only by the producer, but also by neighbouring cells (Buckling et al., 2007; West & Buckling, 2003). *P. aeruginosa* also make a secondary siderophore, pyochelin, which is cheaper to produce but has a lower affinity for iron (Dumas et al., 2013). In contrast, the Pseudomonas heme uptake system (*phu*) is private, as the iron-rich compound heme is taken up directly without the secretion of exploitable exoproducts (Table 1). Additional mechanisms of uptake include the private ferrous iron transport system (*feo*; Table 1 (Cartron et al., 2006)), and the heme assimilation system *(has)*. The *has* system produces a hemophore that can bind to heme, or hemoglobin that contains heme. In the closely related system of *Serratia marcescens* it is conditionally cooperative, as uptake is direct without the hemophore at high concentrations (Ghigo et al., 1997; Létoffé et al., 2004).

*P. aeruginosa* iron metabolism evolves during long term infection. Pyoverdine production has repeatedly been found to be lost during CF infection (Jeukens et al., 2014; Konings et al., 2013; Martin et al., 2011), and the *phu* system has been observed to be upregulated late in infection (Marvig et al., 2014; Nguyen et al., 2014). A major challenge is to identify the selective pressures experienced *in situ* that cause such changes. Host adaptation, to accommodate e.g. antibiotic pressure and resource limitations, is frequently predicted to be the primary force in pathogen evolution (Lieberman et al., 2011; Young et al., 2012). Experimentally recreating host conditions *in vitro* is challenging, given the complex interplay between spatial structure, host immunity and nutrient availability (Clevers et al., 2016), which we are only beginning to be able to measure (Koo et al., 2011). Clinical measurements may differ from experimental *in vivo* and *in vitro* findings (Cornforth et al., 2018), and studies using “simple” experimental conditions do not always give comparable results to those that use more complex, and potentially more clinically relevant conditions (Harrison et al., 2017). In contrast, longitudinal sampling of patients to track *in situ* evolution provides an opportunity to infer selection without making *a priori* assumptions about the experienced environment. Evolutionary theory lets us make testable predictions to distinguish the effect of host adaptations and social interactions. With this approach we have shown that cheating is a major selective force in the loss of cooperative iron acquisition in the CF lung (Andersen et al., 2015).

Here, we identified changes in iron uptake strategies from genome sequences and correlated these with the social environment (the *P. aeruginosa* clonal lineage infecting a host), and duration of infection. The isolate collection covers 546 whole genome sequenced isolates from 61 Danish patients, of 55 clone types (Andersen et al., 2015; Markussen et al., 2014; Marvig et al., 2013, 2015); allowing for the identification of patterns of convergent evolution. While the *P. aeruginosa* populations within patients are remarkably uniform at the clone type level, colonizers diversify during infection (Mowat et al., 2011; Winstanley et al., 2016), and lineages may occupy distinct niches at late stages (Jorth et al., 2015). The isolate collection has been shown to represent the genetic diversity well, with multiple genotypes co-occurring within individual patient samples (Sommer et al., 2016). A key step in our analysis is to categorise isolates with respect to their social environment, for example, presence vs absence of cooperators (pyoverdine producers). This is necessary for testing our hypotheses that iron acquisition strategy is influenced by the social environment. While it is likely that we do not capture all co-infecting strains in our sample, the effect of mis-classifying isolates will homogenise the sample groups and hence obscure rather than amplify any differences, reducing our ability to detect an effect.

**Table 1:**
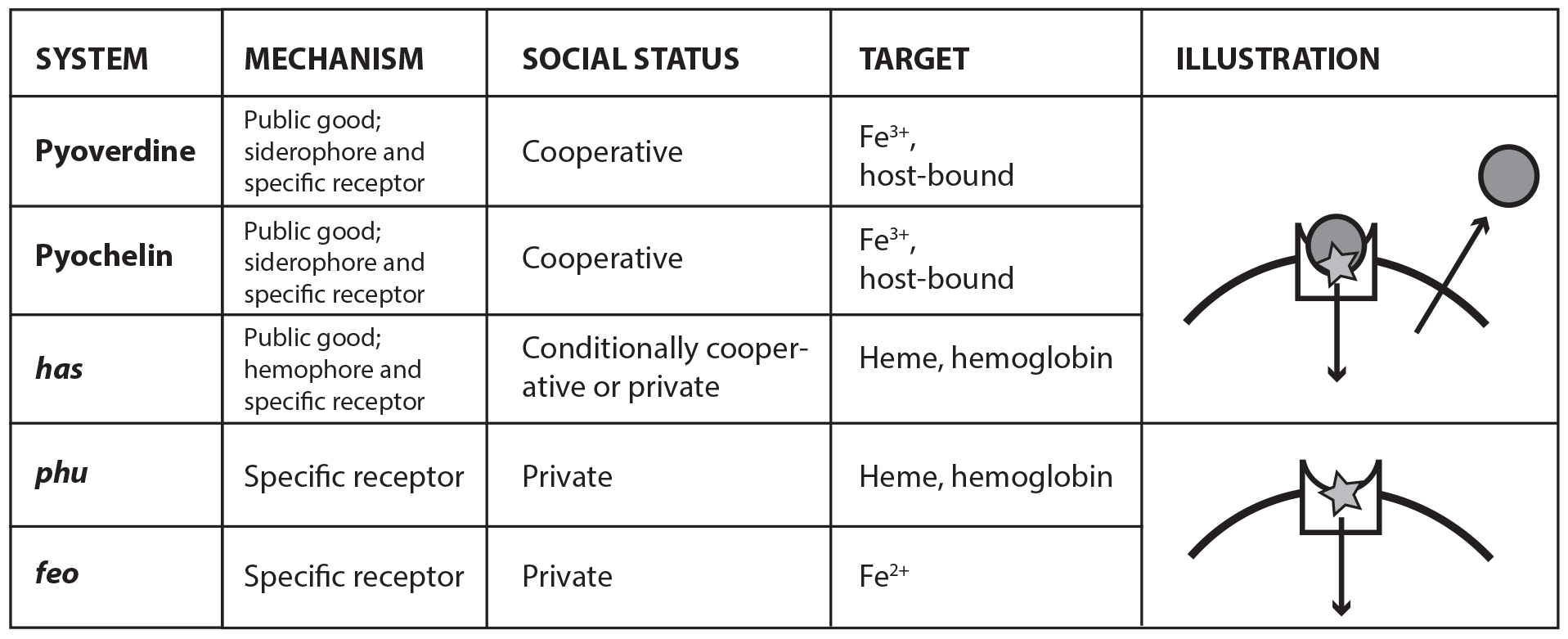
Main iron uptake systems of *P. aeruginosa*. The pyoverdine and pyochelin systems produce siderophores which can steal Fe^3+^ from host carriers. The conditionally cooperative has system produces a hemophore that can bind to heme, or hemoglobin that contains heme. The phu and feo systems are private. All systems have a specific receptor and uptake feeds back to increase system expression. The illustration shows a receptor in the cell membrane taking up an iron source (star). In a cooperative system a secreted public good (circle) binds the target and is taken up by the receptor, whereas a private system takes up the iron source directly.

## Results and Discussion

### More cooperative populations are more vulnerable to exploitation

The environmental clones that infect CF patients produce pyoverdine (Andersen et al., 2015) but what happens as they transition to life in a host? We compared pyoverdine production between strains isolated from different time points of infection in iron-limited media, in which cells must produce pyoverdine to avoid iron starvation. In some clonal lineages of *P. aeruginosa*, pyoverdine production was maintained for >30 years of CF infection: e.g. of the eight clonal lineages of the transmissible clone type DK1 were followed for >30 years; four maintained pyoverdine production (Andersen et al., 2015 and below). Limited diffusion of secreted pyoverdine, as is likely in viscous CF sputum, impairs the ability of potential cheats to exploit and may contribute to stabilize cooperation in these lineages (Julou et al., 2013; Kümmerli, Griffin, et al., 2009); although there may be significant exchange between spatially distinct microcolonies (Weigert & Kümmerli, 2017).
We found that pyoverdine production increased in 33% of clonal lineages within the first two years of infection. Overall, these “super-cooperator” isolates, which had > 30% higher production than isolates initiating infection, were detected in 24 of 45 independent clonal lineages (53%; Fig. 1), and on average sampled 1.86 years into the infection. This suggests that upregulation is initially favoured, likely by iron limitation or inter-species competition. In CF patients, *P. aeruginosa* frequently co-infects with *Staphylococcus aureus* (Folkesson et al., 2012), and co-culture *in vitro*causes an upregulation of pyoverdine production (Harrison et al., 2008). When in co-infection, iron sequestering can have the additional benefit of making it unavailable for *S. aureus* or other competitors (Leinweber et al., 2018; Niehus et al., 2017). Increased production could also be part of a general acute infection phenotype (Coggan & Wolfgang, 2012). For six out of the 24 lineages with super-cooperators we identified candidate mutations in global regulatory and quorum sensing genes that likely cause increased production of pyoverdine as well as other virulence factors, such as pyochelin, pyocyanin and alginate (Table S2). For the remainder, super-cooperators produced pyoverdine at a consistently higher rate than their ancestors but the genetic basis of upregulation was unclear.

Despite the potential benefits of pyoverdine production, the presence of super-cooperators appears to weaken long-term stability of cooperation in the lung. In 12 out of 35 patients, we previously observed the appearance of mutants that have lost the ability to synthesise pyoverdine (Andersen et al., 2015), sampled on average 2.97 years after infection. Here, we found that the likelihood of these non-producers arising was significantly higher in the presence of super-cooperators [χ^2^ (2, N = 47) = 5.7; p < 0.05; fig. 1]. Non-producers were present in four out of 21 clonal lineages without (19%) and in eight out of 24 lineages with super-cooperators (33%).

**Figure 1.**
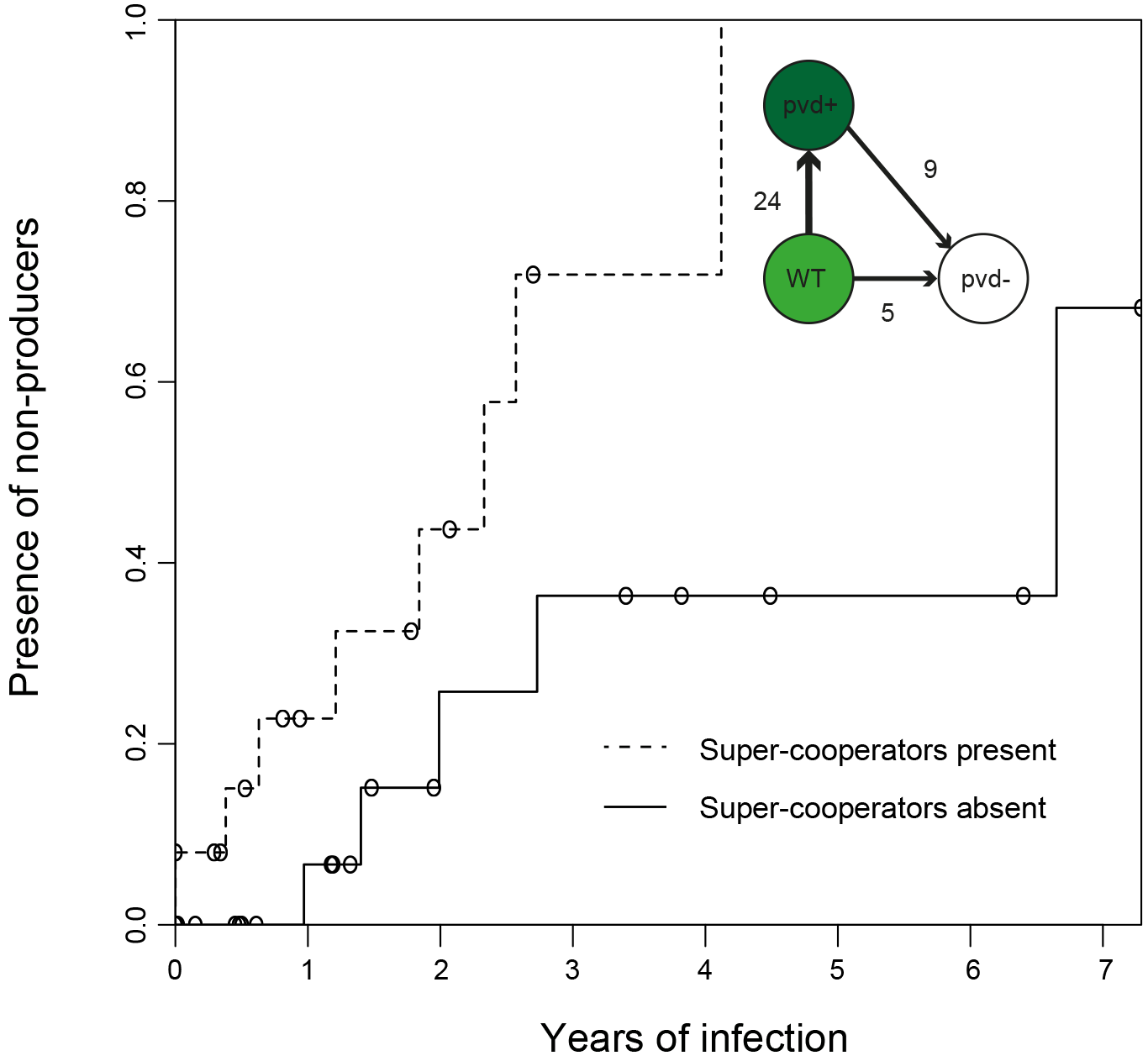
Kaplan-Meier graph showing that pyoverdine non-producers are found more often in the presence of super-cooperators (dotted line, n = 25 representing 24 clonal lineages in patients, one of which had two independent losses of pyoverdine production), compared to when supercooperators are absent (solid line, n = 22, representing 21 clonal lineages in patients, one of which had two independent losses). Circles indicate when sampling of clonal lineages stopped, without the observation of lost pyoverdine production. Insert shows that a transition from wild-type (WT) production to super-cooperation (pvd+) was observed in 24 clonal lineages, followed by nine independent occurrences of non-producers. Non-producers evolved in the absence of supercooperators five times. In total, 33 clone types infecting 35 patients were followed.

Why is upregulation of pyoverdine production associated with the subsequent loss of this trait? Non-producers appear to be favoured as a result of exploiting the pyoverdine production of neighbouring cells - a strategy referred to as cheating (Ghoul et al., 2013). We have previously shown that non-producers in the lung acquire mutations in a pattern consistent with a cheating strategy, as opposed to one expected from redundancy, in that they retained the pyoverdine receptor only if producers were present (Andersen et al., 2015). In other words, non-producers lose the capacity to contribute but retain the ability to benefit from pyoverdine, as long as it is being supplied by cooperative neighbours. The relative fitness of cheats is predicted by theory to be greater when there are higher levels of cooperation in a population, because their competitors are bearing a higher cost (Ross-Gillespie et al., 2007); and this prediction has been shown to hold in empirical tests (Ghoul et al., 2014; Harrison et al., 2008; Jiricny et al., 2010). Hence, supercooperators are predicted to be especially vulnerable to exploitation and invasion of cheats. The pattern of upregulation, followed by loss, is likely to be applicable to the evolution of other exoproducts during infection.

### Evolution of iron acquisition systems in the lung - the sequence of events

How do the changes in cooperative pyoverdine production relate to the overall iron metabolism of the cells? Comparison of the accumulation of mutations across five different systems (Table 1) showed that loss of pyoverdine production is the first change in iron metabolism observed in most clonal lineages. When weighed by the high mutation rate of the system, reflecting a strong selection pressure, and its large size, this is not unexpected [P(X ≥ 16) ~ pois (X; 14.69) > 0.05), fig. 2, table S3, SI text]. We observe cheats appear 14 times without going to fixation during the sampling period, but further record nine cases where no producers are sampled a year prior to, or after emergence a non-producer. One of these is in a transmitted clone type that is subsequently sampled from 16 patients over 35 years. This is the “game over” scenario of cheat invasion. However, because we have continued to sample after this event, we can observe how the population responds to this potentially catastrophic situation. And because cheat invasion is not inevitable, we can compare dynamics in lungs with and without availability of pyoverdine.

### Cheat invasion is associated with a subsequent switch to private iron uptake mechanism

After cooperation was lost, and only if it was lost, we observed that iron uptake was privatised (Fig. 3). This was achieved by upregulation of the private *phu* system. The *phu* system targets the iron-rich heme molecule, which is taken up directly through a membrane-bound receptor (Table 1) (Ochsner et al., 2000). Increased expression of the *phu* system results from intergenic mutations between the *phuS* gene encoding a heme traffic protein, and the receptor gene *phuR* (*phuS*//*phuR* mutations; Marvig et al., 2014). In the isolate collection, a total of 29 SNPs and two indels accumulate in the *phuS//phuR* region. Eight of these mutations in the *phuR* promoter have been found to cause a significant upregulation of promoter activity, and one of these has further been shown, in an isogenic pair, to provide a growth benefit to the carrier when heme is available as an iron source (Marvig et al., 2014) (see also below and Fig. S1). These specific mutations were found to occur significantly more frequently than expected by chance following the loss of pyoverdine production [P(X ≥ 5) ~ pois (X; 0.39) < 0.01); fig. 2]. In three patients, non-producing isolates with *phuS*//*phuR* mutations were found to co-occur with pyoverdine producers, however in two of these the producer and non-producers were unlikely to interact (SI text).

**Figure 2:**
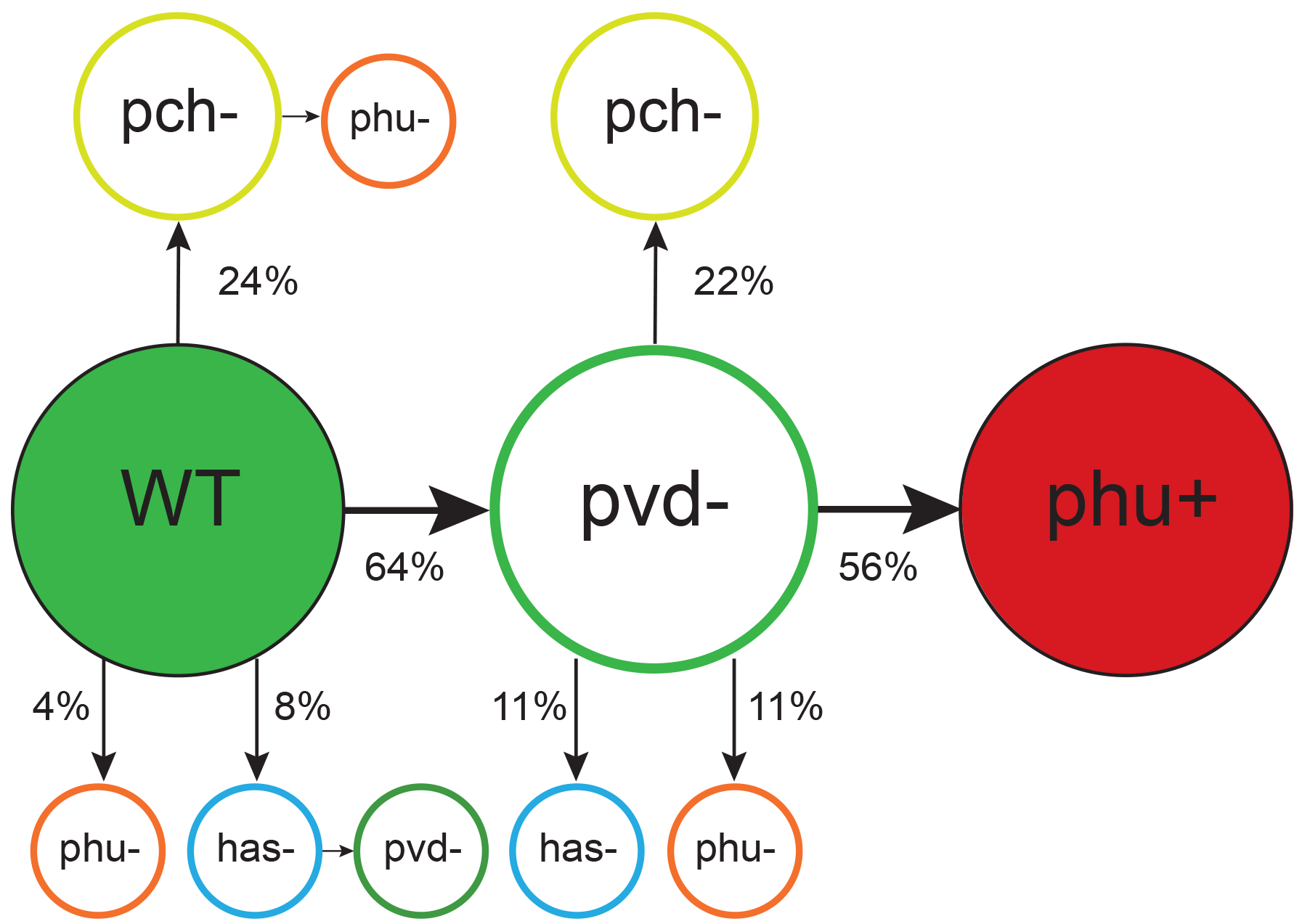
Order of mutations across iron acquisition systems. Wild type-like (WT) isolates colonize patients, and subsequent loss of cooperative pyoverdine production (pvd-) is the most common change in iron metabolism (n = 25), compared to mutations expected to affect the pyochelin, phu and has systems negatively (pch-, phu-, has-). Loss of pyoverdine production is followed by intergenic phuS//phuR mutations, predicted to upregulate the private phu system, significantly more often than expected by chance (phu+, n = 9). Colour indicates iron uptake system (green = pyoverdine, yellow = pyochelin, dark red= intergenic phuS//phuR, orange = phu (either a mix of phuS//phuR and other phu mutations, or only ns SNPs), blue = has). The occurrence of has mutations may be underestimated due to low sequencing depth of two genes. The figure shows only transitions where there was a clear order in which systems were affected, see table S3 for all transitions.

Social interactions are unlikely to be the only selection pressure influencing dynamics in iron acquisition strategies. Changes in iron uptake strategy might more readily be expected to reflect changes in the lung environment as the availability of iron in different forms increases with disease progression (Hunter et al., 2013). And while iron is found to be equally available in its oxidized ferric (Fe^3+^) and reduced ferrous forms (Fe^2+^) in early infection, there is a slight skew towards the reduced form later, which the *feo* system targets (Hunter et al., 2013). Heme and hemoglobin are also present in the sputum (Ghio et al., 2013), and availability has been speculated to increase as the lung tissue degrades (Marvig et al., 2014), as evidenced by increased coughing up of blood, termed hemoptysis, with age (Thompson et al., 2015). As such, pyoverdine may be most efficient early in infection, and *phu* only advantageous later as lung tissue degrades and/or alternative sources of iron become more or less abundant. However, the patterns we observe are not consistent with this explanation. If pyoverdine production is maintained, the *phu* genotype remains unaltered from colonization, even after 35 years of infection. The*phuS//phuR* mutations, responsible for upregulating the private *phu* system, are only observed in non-producers [χ^2^ (2, N = 64) = 12.1; p < 0.01; fig. 3 & 4], occurring between 1.1 to 35 years after the loss of pyoverdine production. Upregulation of private iron uptake can, therefore, be predicted from the social environment, not duration of infection and associated changes in the lung environment.

**Figure 3:**
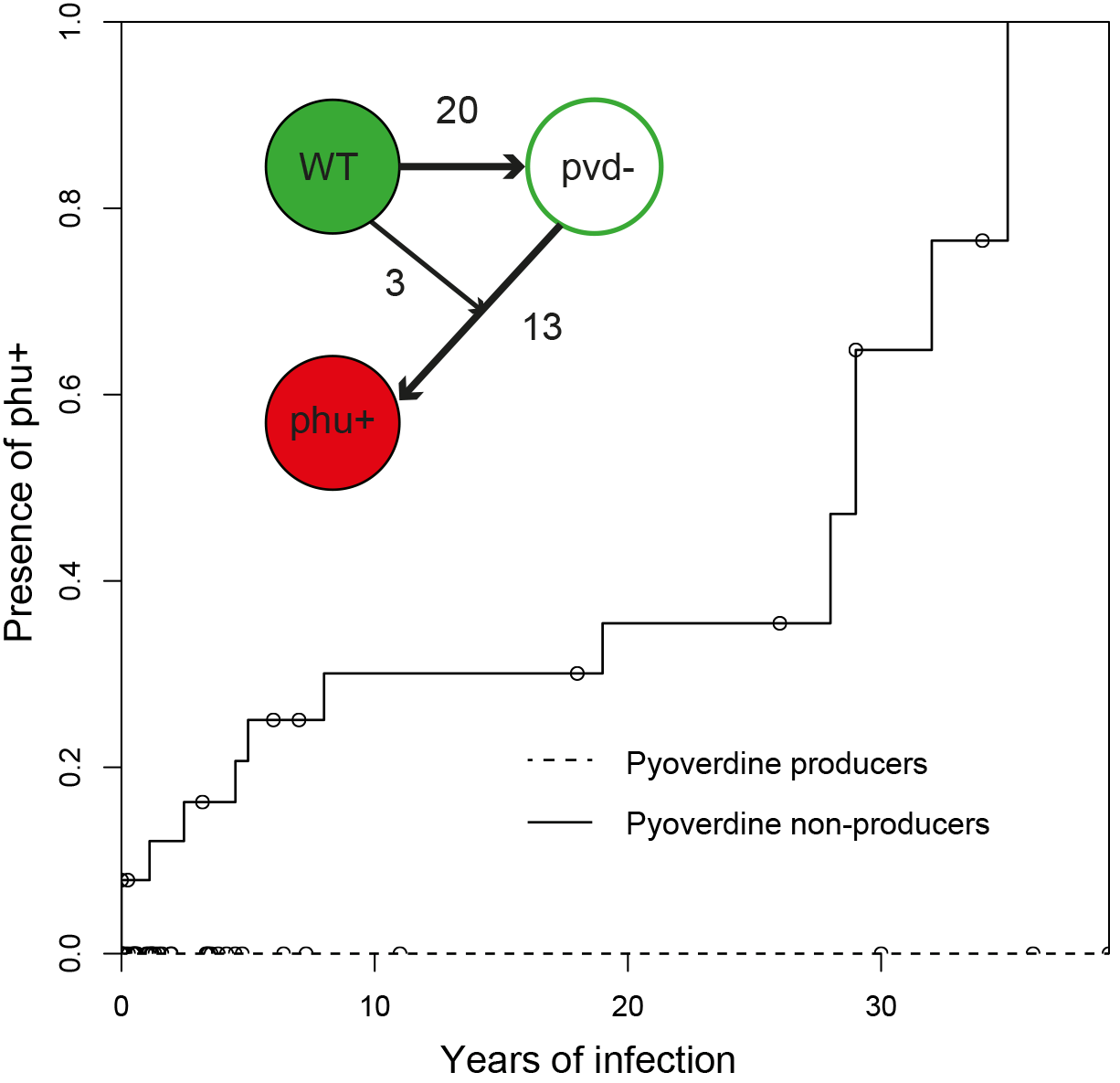
Kaplan-Meier graph showing that phuS//phuR mutations only are found in clonal lineages that do not produce pyoverdine (solid line, n = 26), and never when pyoverdine is produced (dashed line, n = 38). Circles indicate when sampling of clonal lineages stopped, without the observation of phuS//phuR mutations. Insert shows that a transition from wild-type pyoverdine production (WT) to no production (pvd-) was observed 23 times in clonal lineages, while phuS//phuR mutations occurred 16 times independently (phu+), 13 of which were after pvd-mutations. The order could not be inferred in three cases.

**Figure 4:**
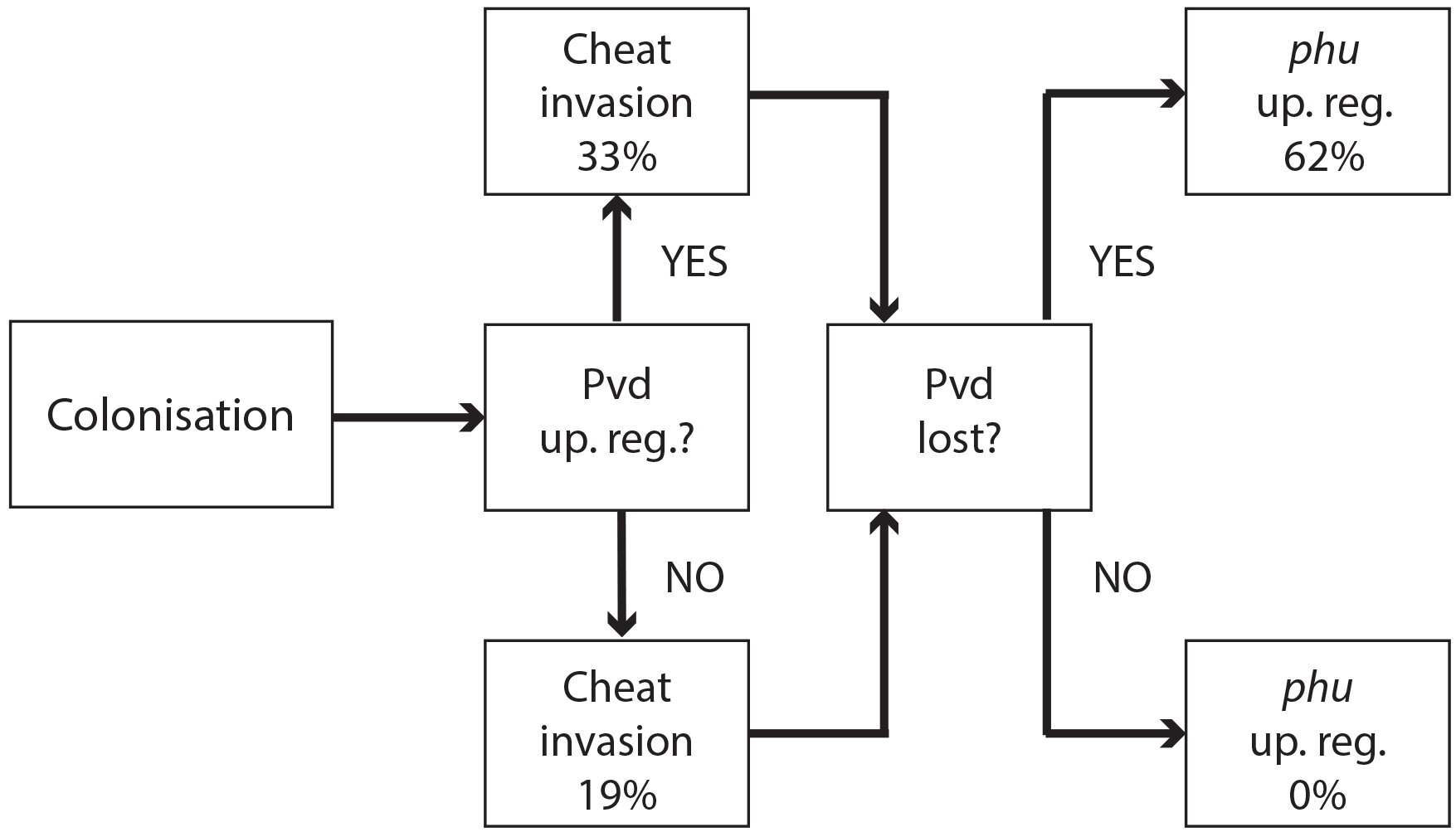
Evolutionary trajectories of iron metabolism in the CF airways. Following infection (n = 45 clonal lineages of 33 clone types), pyoverdine production is frequently upregulated (n = 24). Cheaters are significantly more likely to invade following upregulation. Following loss of pyoverdine production (n = 26 non-producing clonal lineages of 7 clone types, from 23 independent losses of production), the phu system is frequently upregulated by intergenic phuS//phuR mutations (n = 16), while this never occurs when pyoverdine is produced (n = 38 producing clonal lineages of 30 clone types).

### Why is private heme uptake only upregulated if cooperation is lost?

The fact that we only observe upregulation of the *phu* system if pyoverdine production has been lost suggests that privatisation is not universally beneficial but may instead be a solution of “last resort” when other mechanisms are no longer functioning. *P. aeruginosa* iron uptake is finely controlled by feedback loops, in response to environmental iron concentrations and need (Vasil & Ochsner, 1999). As iron is toxic at high concentrations (Anzaldi & Skaar, 2010; Touati, 2000), an indiscriminate upregulation of one system could be a significant disadvantage in environments where iron concentrations fluctuate. We, therefore, tested whether the growth benefits of *phu* upregulation (by *phuS//phuR* mutations) were dependent on iron availability, in the form of heme. We examined the growth difference between a clinical isolate that had lost pyoverdine production, and an isolate, isogenic but for a clinical *phuS//phuR* mutation that cause an upregulation of *phuR*(Marvig et al., 2014). The experiment was performed in iron-limited media supplemented with heme at biologically relevant concentrations, which ranged from < 1 μM typical of healthy individuals, across 2.5μM, 5μM to *ca.* 10 μM typical of CF patients (Ghio et al., 2013).

Consistent with our hypothesis, we find that *phu* upregulation is only beneficial at a narrow range of intermediate heme concentrations. There was a significant growth difference between the isolates and heme concentrations, and an interaction effect (Two-Way ANOVA; Interaction term: Isolate*Heme: F = 48.11, df = 3, p < 0.01; fig. 5; table S4). At 2.5 μM heme the isolate with an upregulated*phu* system achieved a higher density (Tukey HSD, p_adj_ < 0.01, fig. 5), consistent with previous findings (Marvig et al., 2014). In contrast, at 5 and 10 μM the isolate without upregulation had an advantage (p_adj_ < 0.05, Fig. 5). No growth was observed in the absence of heme, and at 1 μM there was no significant difference in growth between the isolates (p_adj_ = 0.073). This suggests that *phuS//phuR* mutations lead to increased heme uptake that is beneficial at low external concentrations (>1 and < 5 μM heme) but detrimental at high concentrations (> 5 μM heme). If iron availability increases during infection, an initially beneficial upregulation of heme uptake would turn toxic. In the *phu* system we found a significant bias towards mutations of the *phu* receptor gene *phuR* (P(X ≥ 24) ~ pois (X; 10.69) < 0.05; table S5, SI). These were not randomly distributed but primarily found in the extracellular loops of the *phu* receptor, that initiate the uptake of heme (Noinaj et al., 2010; Smith et al., 2015)[P(X ≥ 15) ~ pois (X; 7.28) < 0.05; fig. S2]. If there is selection to reduce iron concentration within cells, these may, therefore, represent compensatory mutations.

**Figure 5:**
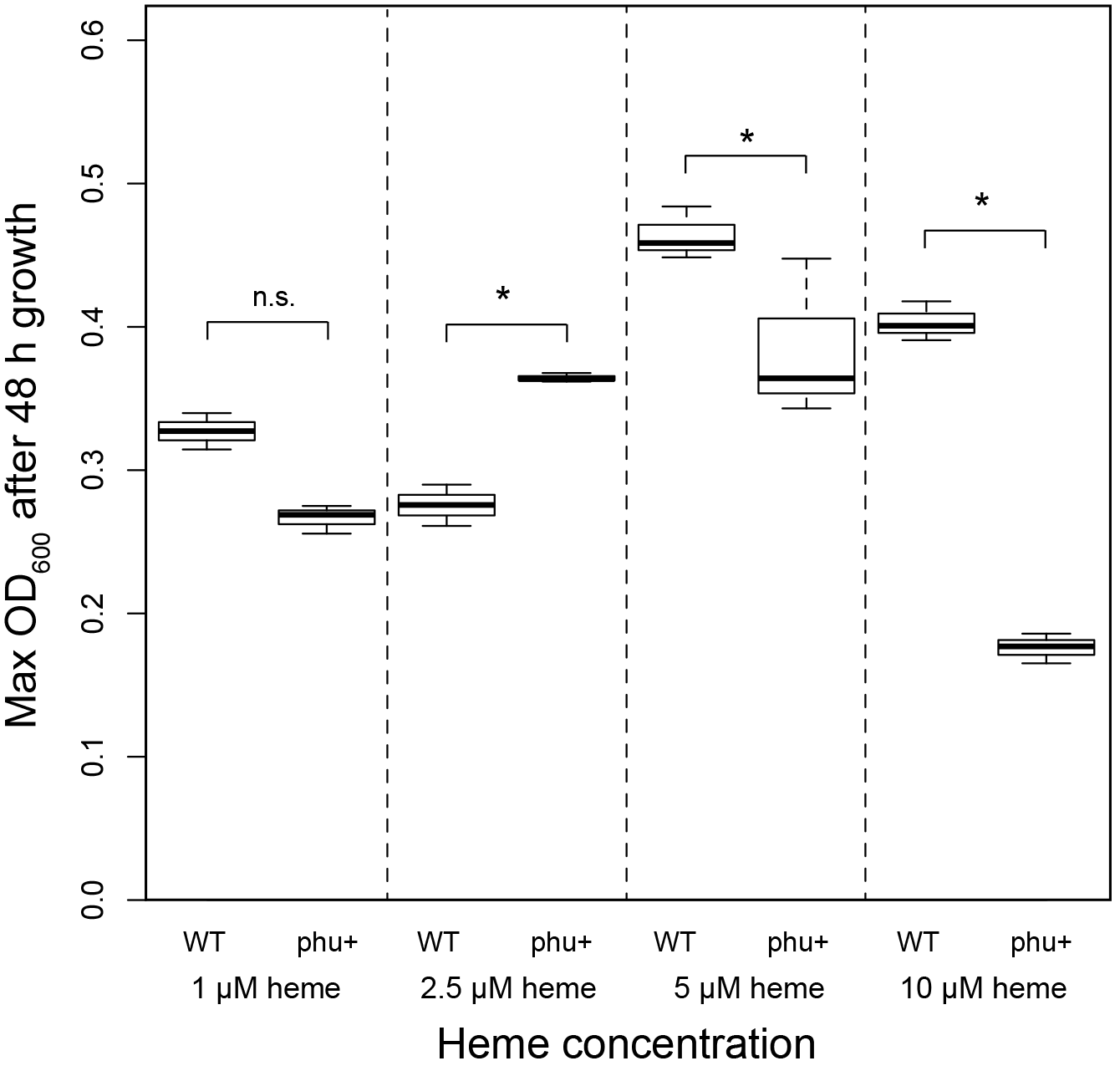
Effect of phu upregulation depends on the heme concentration. An isogenic pair of isolates differing only in a 1 bp phuS//phuR deletion was grown in iron-limited media supplemented with heme at 1-10 μM. The clinical isolate without the mutation, that does not produce pyoverdine (WT), had a significant higher maximum OD_600_ after 48 h growth at 5 and 10 μM, whereas the mutant (phu+) grew best at 2.5 μM. There was no significant difference at 1 μM. Boxplots show median value, and the interquartile range ±25%. The * marks a significant difference at p < 0.05.

## Conclusions

Bacterial cells will be under selection to adapt to the host environment, including antibiotic pressure (Lieberman et al., 2011). The observations we report here, however, demonstrate the errors of interpretation that can arise from assuming that bacterial cells are driven entirely by the evolutionary imperative to survive in the host. As in all other organisms, fitness is also determined by the ability to compete with members of the same species, and these two evolutionary drivers - survival and competition - do not always coincide. The effect of the social environment on iron acquisition mechanism remains statistically detectable despite the noise from variation in patient age, sex, infection and treatment. In a clinical system like this, failure to appreciate these factors can lead to misinterpretation of how bacterial cells are influenced by the host environment and, therefore, impair necessary understanding for the development of intervention strategies (Leggett et al., 2014).

Evolutionary theory clearly predicts that any adaptation involving cooperation can be vulnerable to exploitation, irrespective of its benefits (Axelrod & Hamilton, 1981). But this is often overlooked when it is difficult to characterize the selective environment, as for example, inside a human host. By longitudinal sampling of bacterial populations, we were able to observe the consequences of cheat invasion for an essential trait and show that switching to a private strategy can provide bacterial cells with an evolutionary “escape route”. This observation highlights the importance of considering potential constraints imposed by social interactions: if a private trait is expressed it may not be the most efficient in a given environment, but simply the “last resort” in a population with a history of exploitation.

## Acknowledgements

We thank Stuart West, Lars Jelsbak and Rolf Kümmerli for comments on the manuscript. Trine Markussen and Lars Jelsbak provided analysed sequences of additional DK1 isolates, and Hossein Khademi provided valuable discussion of the *phu* mutations. This work was supported by grants from Villum Fonden (grant 95-300-13894) and Lundbeckfonden (grant R108-2012-10094) to SBA, the European Research Council (grant SESE) to ASG, and Rigshospitalet (grant R88-A3537), Novo Nordisk Fonden (grant NNF12OC1015920) and Lundbeckfonden (grant R167-2013-15229) to HKJ.

## Author contributions

SBA & ASG designed the study, SBA & RLM analysed the data, MG & ZBL performed the experimental work, HKJ and SM acquired the isolates and all related information. All authors contributed to the interpretation of the data, the writing of the manuscript, and approved the final version.

## Ethics statement

The CF isolates used for sequencing were obtained from the Department of Clinical Microbiology, Rigshospitalet, Copenhagen, where they had been isolated from CF patients treated at the Copenhagen CF clinic. The use of isolates was approved by the local ethics committee at the Capital Region of Denmark Region Hovedstaden: registration number H-1-2013-032. All patients have given informed consent. In patients under 18 years of age, informed consent was obtained from their parents. All patient isolates were anonymized prior to any experimental use or analysis.

## Methods

### Isolate collections

Two collections of whole genome sequenced *P. aeruginosa* isolates were used, as described previously (Andersen et al., 2015). Of the transmissible clonetypes DK1 and DK2, 95 isolates from 25 CF patients were sampled between 1973 and 2012. (Jelsbak et al., 2007; Markussen et al., 2014; Yang et al., 2011) An additional 10 DK1 isolates were included compared to Andersen et al. (2015). Further, 451 isolates of 54 clonetypes from 36 young CF patients were used (the “children’s collection” (Andersen et al., 2015; Marvig et al., 2015), of which 10 were of the DK1 clone type). The sampling regime for each patient is described in Table S6. The two collections complement each other as the first covers a long time period, with relatively few samples from each patient and of both early and long-term infection stages, whilst the second covers the first 10 years of infection with more extensive sampling within patients.

The length of infection of each clone type in each patient was calculated. Clone types sampled only once from a patient were excluded. For the transmissible clone types DK1 and DK2 the total length of infection across patients was used (39 and 35 years, respectively). This is a very conservative estimate of length of infection, as evolution occurs independently in different patients. To calculate the number of generations, a doubling time of 50 min. was used for clonal lineages infecting for less than 10 years, and for the DK1 and DK2 isolates a doubling time of 65 min. based on the measurements of Yang et al. (2011) (Table S1).

### Changes in pyoverdine production during infection

We tested if loss of pyoverdine production is correlated with preceding upregulation. Measurement of pyoverdine production was carried out by fluorescence readings at 400/460 nm excitation/emission with a 475 nm cut-off following growth in iron-limited media (Andersen et al., 2015; Kümmerli, Jiricny, et al., 2009). For each clonal lineage the baseline pyoverdine production was determined as that of the first isolate(s) of a clone type from a patient, and the production of all subsequent isolates of this clone type was compared to this. An isolate was classified as an overproducer if its production was > 30% higher than that of the first. This cut-off was chosen as ~95% of the isolates had a lower standard deviation between replicates. Loss of pyoverdine production was defined as sampling of a non-producing isolate from a clonal lineage that produced pyoverdine at the beginning of the infection period. The analysis was performed on longitudinally sampled clone types from the children’s collection only, as the DK1 and DK2 isolates were not sampled frequently within patients. In all but three cases, over-producers were sampled before non-producers. In two cases there was < 5 months between sampling, and we scored non-producers as occurring at the same time as overproducers. In one case the non-producer was sampled 2.95 years prior to the over-producer and scored as having occurred in the absence of an over-producer. In two clone types two independent transitions to non-producers were observed and these were included as independent events.

We categorized each clone type in individual patients as harbouring over-producers or not. If an over-producer was present, the length of time to a non-producer was observed, or till the last sample was collected from the patient, was noted. Clone types without non-producers were classified as right-censored. The same was done for patients without over-producers. In total, 45 clonal lineages were followed (some clone types occur in multiple patients, and some patients are infected by multiple clone types (Marvig et al., 2015)). Clone types DK1, DK11, DK40 and DK53 were excluded from the analysis, as they were “chronic” clone types with no pyoverdine producers sampled (Andersen et al., 2015). The timing of sampling of non-producers was analysed with the “survival” package in R (R Core Team, 2013; Therneau, 2015).

### *Identification of mutations in the pyoverdine, pyochelin,* phu, has *and* feo *systems*

Mutations in the pyoverdine (pvdQ-pvdI + pvdS-pvdG + pvcA-ptxR + fpvB), the pyochelin (*ampP-pchA*), the *phu* (*prrF1-phuR*), the *has* (*PA3404-hasI*) and the feo operons (*PA4357-feoA*) were identified previously from Illumina GAIIx or HiSeq2000 reads (Andersen et al., 2015; Markussen et al., 2014; Marvig et al., 2013, 2015; Yang et al., 2011) (Table S5, S7). Mutations of iron systems in the 10 new DK1 isolates were identified as described previously (Andersen et al., 2015). Mapping of reads revealed low sequencing depth of the *hasD* (required for hemophore secretion) and *hasS* (the *has* anti-sigma factor) genes, likely due to repetitive regions. These were unsuccessfully attempted amplified by PCR. Mutations were categorized as synonymous (syn) or non-synonymous (ns) SNPs, insertions or deletions (indels), or intergenic. We tested for a bias in which genes in each operon were mutated with [P(X ≥ changes_observed_) ~ Poisson distribution (pois) (X; changes_expected_) < 0.05].

We identified the order of mutation across the different systems, with a system classified as mutated by the presence of ns SNPs or indels in any gene in the system. For pyoverdine, the presence of mutations was compared to the actual measurements of production; the two measures are highly consistent (Andersen et al., 2015) but when incongruent the phenotype was used. For the *phu* region intergenic *phuS//phuR* mutations predicted to cause receptor upregulation were included, and when occurring alone these were recorded separately. We calculated the expected order of mutation, equally weighted by the size and number of mutations of the different systems. We tested, as described above, whether this was different from the observed, using only cases where there is a clear order of mutations (Fig. S1).

For the *phu* system, we located mutations in the *phuR* receptor gene to functional regions (the N-terminal-domain, periplasmic turns, transmembrane strands or extracellular loops) using a predicted structure of the receptor (http://bioinformatics.biol.uoa.gr//PRED-TMBB (Bagos et al., 2004)). For each domain in the receptor we calculated the expected number of non-synonymous amino acid changes, as the proportion the domain comprises of the entire protein times the observed number of changes. Whether the observed changes in each domain differed significantly from the expected was calculated as described above.

### Transitions between public and private iron uptake

To test if the appearance of intergenic *phuS*//*phuR* mutations was correlated with loss of pyoverdine production we categorized each independent acquisition of these mutations as having occurred in the presence or absence of measured pyoverdine production. In the absence of pyoverdine production the time between loss of production and occurrence of mutations was estimated. The analysis was performed on longitudinally sampled clonal lineages from both collections of isolates, that is lineages with at least two isolates with or without pyoverdine production. Further, three cases where the loss of pyoverdine production and *phuS//phuR* mutations were observed in the same isolate were also included. Clonal lineages that did not acquire *phuS*//*phuR* mutations were classified as right-censored. For DK1 and DK2 we used monophyletic clades with and without mutations as independent clonal lineages. We included isolates of DK53 where no WT pyoverdine producer was sampled, but *phuS//phuR* mutations were inferred by comparison to PA01 (Table S5). The timing of sampling of mutants was analysed with the “survival” package in R (Therneau, 2015). We are likely to over-estimate the length of time to mutation, as patients chronically infected with the transmissible DK1 and DK2 clone types were infrequently sampled (Markussen et al., 2014; Marvig et al., 2013). In these cases, with several years between longitudinal samples, a mutation may only be detected years after occurring.

### *Uptake of heme following* phuS//phuR *mutations*

We tested if clinical isolates with *phuS//phuR* mutations have a growth advantage in iron-limited media supplemented with heme, when matched to their closest genetically related isolate without mutations. The *phuS//phuR* mutations occurred in three different clone types without additional *phu*mutations (DK1, DK2 and DK32, highlighted in yellow in Table S5). The six isolates with mutations were matched with four close relatives without, because DK2 in two instances had independent *phuS//phuR* mutations in two closely related isolates. One additional DK2 isolate with a mutation was lost from the collection, and only available as sequence reads (P80F1, table S5). As a positive control we included an isogenic pair of DK2, only differing in the presence of a 1 bp *phuS//phuR* deletion (Marvig et al., 2014). This pair represents the WT isolate used in two of the clinical pairs, and the mutation of one of the clinical isolates moved to this background (P0M30-1979 and P173-2005; Fig. S2). Three biological replicates of each isolate were cultured in liquid LB media for 24-48 h, so that all cultures reached an OD_600_ above 0.1. Cultures were subsequently standardised to a starting OD_600_ of 0.001, in iron-limited CAA media following Kümmerli, Jiricny, et al. (2009), without heme or supplemented with 2.5 or 5μM heme as an iron source (BioXtra porcine hemin ≥ 97.0%, Sigma-Aldrich). Of each culture 200μL were inoculated in 96 well plates. Plates were sealed with a breathable membrane to avoid evaporation and incubated in a plate reader measuring OD_600_ every 30 min for 48h. Growth curves were plotted in Excel and the average maximum OD_600_ of the three replicates calculated. The difference in max OD_600_ for a pair of related isolates with and without *phuS//phuR* mutations was analysed with a t-test in R (R Core Team, 2013). The isogenic control pair was also grown supplemented with 1 and 10μM heme, and the differences in max OD_600_ were compared in R with a Two-Way ANOVA test with post hoc comparisons with Tukey HSD.

## Supplementary text

### Order of mutations across iron uptake systems

Loss-of-function of the cooperative pyoverdine system occurred earliest in 16 out of 25 clonal lineages where the sequence analysis provided a clear order of systems affected (Fig. 2; table S3). These were observed to occur on average 3.21 years after the first sampling of the clone type in the young patients (n = 11 clonal lineages in nine patients), and between 0-39 years after infection for the more rarely sampled transmissible clone types (n = 5 clonal lineages in five patients). Mutation of the pyochelin system first was found not to differ from that expected given a random distribution of mutations [P(X ≥ 6) ~ pois (X; 5.14) > 0.05), fig. 2; table S3].

The conditionally private or cooperative *has* heme uptake system was found to acquire few mutations during infection. It remains to be tested if this is because maintenance of the system is favoured by selection, or if selection for loss is relatively weak. The private *feo* system is rarely mutated (Table S3) and has been suggested to be of increased importance as conditions turn microanaerobic during infection (Hunter et al., 2013).

### *Co-occurrence of isolates with* phuS//phuR *mutations and pyoverdine producers*

The *phuS//phuR* mutations only occurred in isolates that had lost pyoverdine production. In three instances these, however, co-occurred in the patient with isolates that produced pyoverdine. The producers and non-producers with *phu* upregulation are unlikely to have interacted at the time when the iron metabolism mutations occurred in two of these. In patient P30M0, infected with clone type DK1, the cooperating isolates were sampled from the patient’s sinuses only, while the nonproducers with *phu* upregulation came from lung samples. In patient P28F1, two lineages of clone type DK1 co-occurred. One had been transmitted from another patient, where pyoverdine production was lost, and *phu* likely upregulated, prior to transmission and establishment of a coinfection. In both cases the non-producers carried mutations in the pyoverdine receptor. In patient P83M2 a pyoverdine producer and an isolate with a *phuS//phuR* mutation, both of clone type DK32, were found in the same lung sample. The latter harbours the most common *phuS//phuR* mutation (G 146 upstream//35 upstream A, table S5), which is located outside the *phuR* promoter (Marvig et al., 2014) . It is the only isolate to have this *phuS//phuR* mutation in isolation, all others with it have an additional SNP, and the effect of it is unknown (but see below).

### *Effect of* phuS//phuR *mutations in clinical isolates*

A range of the clinical *phuS//phuR* mutations have been shown to cause increased promoter activity, increased expression of the *phuR* gene and a growth benefit in iron-limited media supplemented with heme (Marvig et al., 2014; Yang et al., 2011). We tested if the *phuS//phuR* mutations are also beneficial in the clinical isolates, using six isolates of three different clone types that had acquired a variety of *phuS//phuR* mutations, and no additional *phu* mutations (Table S5). We compared fitness of the mutants with their closest genetically related isolate without *phuS//phuR* mutations, to control for phylogenetic differences. The isolates in a pair were, however, separated by multiple other mutations as they were sampled up to 26 years apart. The pairs were grown in iron-limited media with 2.5 and 5 μM heme added, which corresponds to medium concentrations observed in CF patients (Ghio et al., 2013). Three mutants with intergenic SNPs had significantly higher (P73F1; 26% higher; t = -6.33, df = 3.48, p > 0.01 and P83M2; 16% higher, t = -4.48, df = 2.15, p < 0.05; isolates sampled 0 and 23 years apart) or a tendency towards a higher maximum OD_600_ in 5 μM heme compared to their closest relative (P66F0; 17% higher, t = -2.85, df = 2.67, p = 0.07; isolates sampled 10 years apart; fig. S2). A mutant with a three bp insertion, and an isolate with a one base pair deletion in *phuS//phuR* in addition to a SNP, showed no significant difference in growth compared to their closest relative (17% higher, t = -1.78, df = 2.95, p = 0.18; 16% lower, t = - 0.55, df = 2.56, p = 0.10; isolates sampled 3 and 26 years apart; fig. S2). These two isolates have, however, previously been shown to have respectively 25 and 116 fold increased expression of the *phu* receptor compared to the wild type (Marvig et al., 2014; Yang et al., 2011), and transfer of the one base pair deletion to its ancestor confirms that it infers growth benefits in heme media (Marvig et al., 2014) (fig. 4). This suggest that other mutations across the genome affect the phenotype we are interested in, which is not unexpected given that we are comparing isolates sampled years apart, and the complex genome of *P. aeruginosa* where almost 10% of the genes are predicted to be regulatory (Stover et al., 2000). There were no significant growth differences with 2.5 μM heme added. Both isolates of the sixth pair, with a SNP, grew very slowly in rich LB media, and not at all in iron-limited media with or without heme. Caution is generally required when interpreting results of behavioural assays in the lab as evidence of the phenotype *in situ*, as growth conditions are difficult to replicate *in vitro*. However, this data provides confirmation that the mutations we observe in the lung isolates upregulate the *phu* iron uptake pathway *in vitro*.

### Supplementary figures

**Figure S1:**
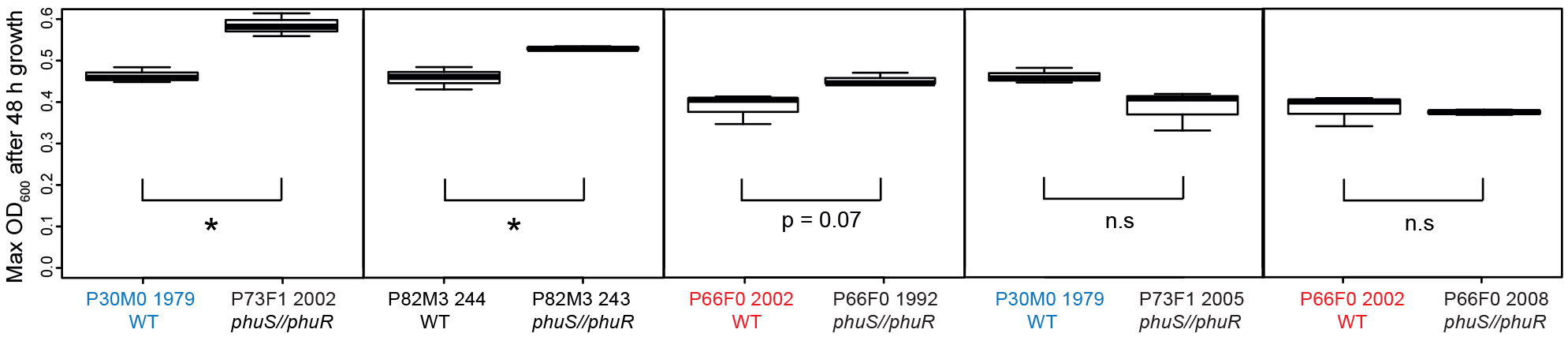
Growth effect of *phuS//phuR* mutations in clinical isolates. Isolates with *phuS//phuR* mutations were grown in iron-limited media supplemented with 5μM heme for 48 h and compared with their closest relative (WT). Two isolates with mutations had a significantly higher maximum OD_600_ than their closest WT relative, and one showed a trend. Pairs where the same WT isolate is used because of two independent acquisitions of *phuS//phuR* mutations are highlighted in red and blue. Boxplots show median value, and the interquartile range ± 25%. * marks a significant difference.

**Fig. S2:**
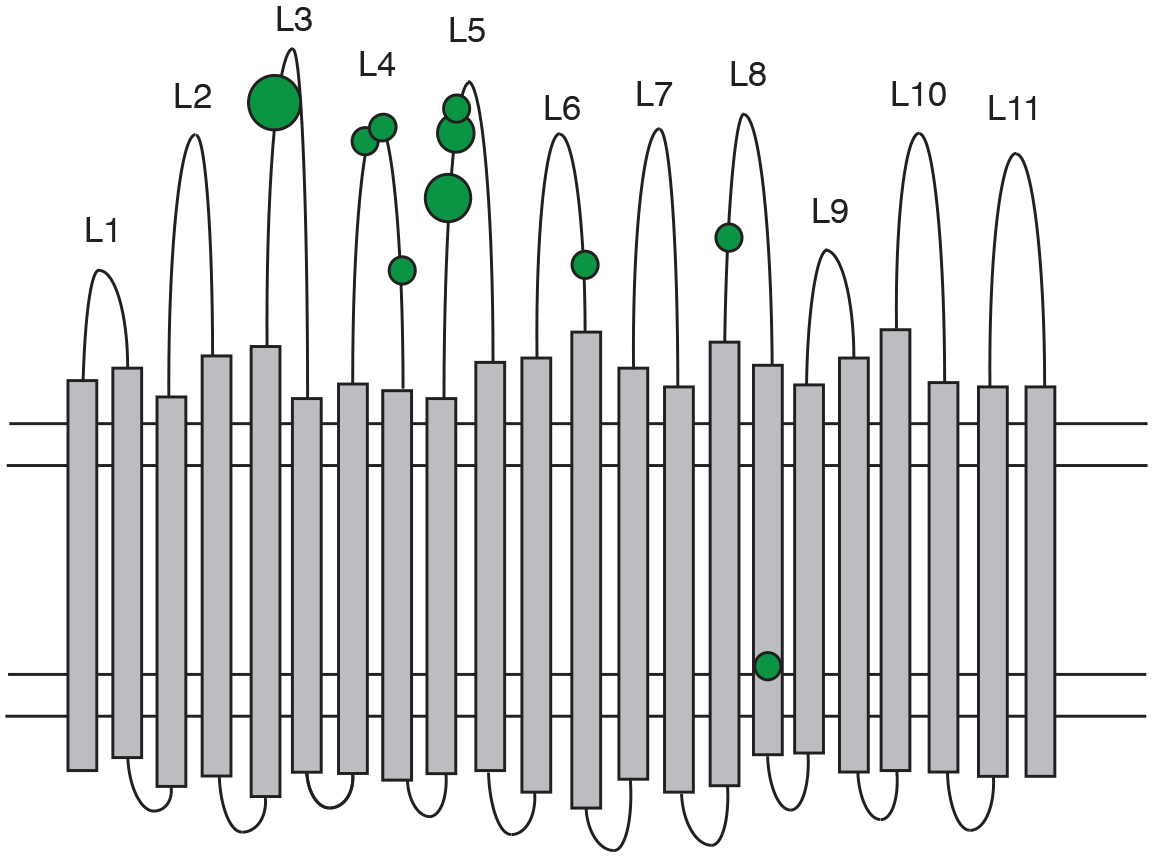
Distribution of non-synonymous mutations in the phuR heme receptor. The figure shows the β-barrel structure spanning the cell membrane (horizontal lines) with transmembrane strands (grey bars), extracellular loops (labelled L1-L11) and periplasmic turns (bottom loops). Green circles indicate the location of non-synonymous SNPs, the smallest circles denoting one mutation and the largest four. Mutations were significantly biased towards the extracellular loops.

### Supplementary Tables

**Table S1:** Calculation of estimated generation times of infecting clonal lineages. For each, the patient ID, clone type, length of infection in years and minutes, estimated doubling time and calculated number of generations is given.

**Table S2:** Mutations identified in pyoverdine over-producers that may contribute to this phenotype. For each mutation, the patient ID, clone type and gene is listed, as well as a reference to previous work showing a link between the gene and pyoverdine production.

| Patient ID | Clone type | Mutation | Reference |
| --- | --- | --- | --- |
| P40M5 | DK43 | ns SNP in PA2384 | (Ochsner et al., 2002) |
| P67M4 | DK41 | Mutations in *gacA*, *gacS* and *retS* | (Frangipani et al., 2014) |
| P82M3 | DK32 | Insertion in *mucA* | (Wu et al., 2004) |
| P98M3 | DK36 | Deletion in *mucA* | (Wu et al., 2004) |
| P99F4 | DK06 | ns SNP in *lasR* | (Stintzi et al., 1998)<br>(negative effect on<br>production in knock-out) |
| P19F5 | DK15 | ns SNP in *mucA* | (Wu et al., 2004) |
| P50F5 | DK07 | Insertion *wspR* | (Visaggio et al., 2015) |

**Table S3:**
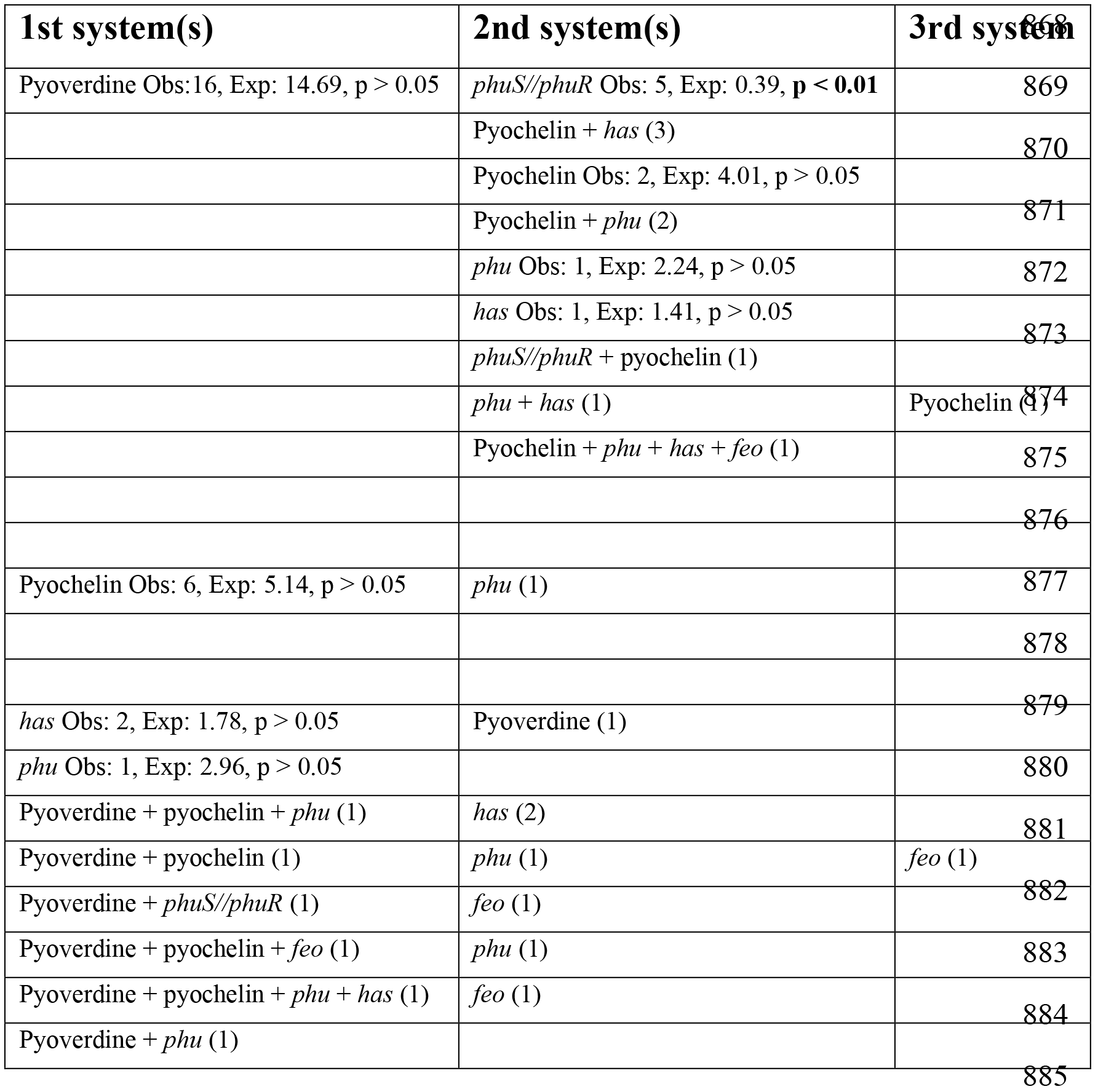
Transitions between iron uptake systems characterized as the order in which mutations accumulate. When a clear order of mutations could be established the observed and expected number of transitions, given a Poisson distribution, is stated and whether the difference between them is statistically significant.

**Table S4:** Results of Tukey HSD post hoc comparison of a Two-Way ANOVA analyzing the difference in Max OD_600_ between an isogenic pair of isolates differing in a *phuS//phuR* mutation at four different heme concentrations. (Excel sheet)

**Table S5:** List of mutations in genes associated with iron metabolism. For each isolate, the patient ID, clone type and mutations are listed. A // denotes an intergenic mutation. The isolate names refer to previous descriptions (Andersen et al., 2015; Markussen et al., 2014; Marvig et al., 2015). The ten DK1 isolates that were not included in the previous analysis of pyoverdine mutations had one ns SNP in pvdL (A5702G) and one in pvdP (C1508T, patient P55M3, isolate TM50 & TM51). The six isolates used in the phenotypic assay of *phuS//phuR* mutations are highlighted in yellow. (Excel sheet)

**Table S6:** Overview of patients and isolates. (Excel sheet)

**Table S7:**
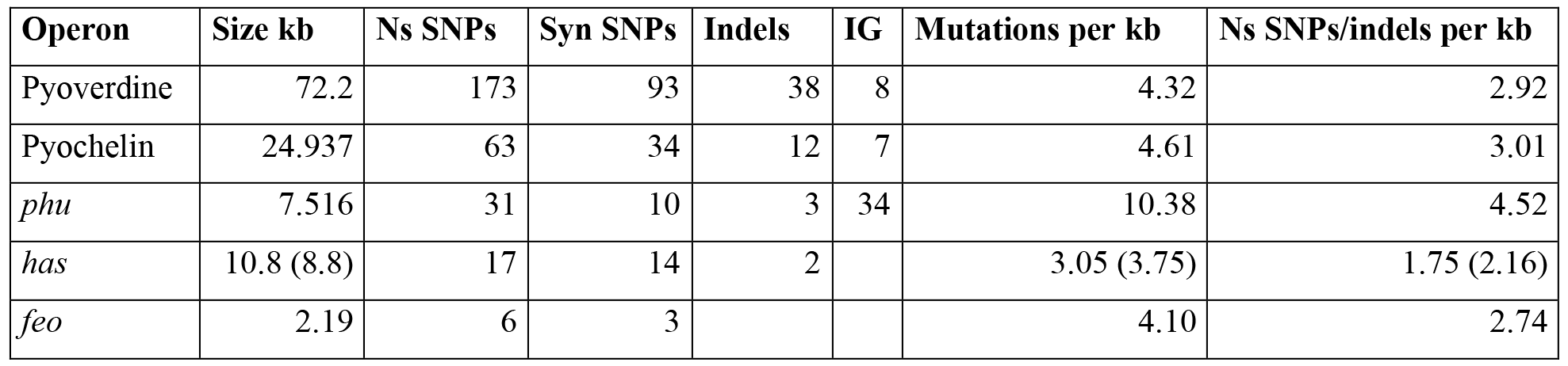
Summary of the mutations found in five different iron uptake systems in *P. aeruginosa*. The size of the respective operons is given in kb, and mutations are listed as non-synonymous (ns) SNPs, synonymous (syn) SNPs, insertions and deletions (indels) and intergenic (IG). The overall mutations, and ns SNPS and indels, per kb are listed. The *phu* operon experienced the highest mutation rate, dominated by intergenic mutations. Two genes in the *has* operon were poorly sequenced in the majority of isolates, and this is taken into account in the size and mutations listed in parentheses.

| Operon | Size kb | Ns SNPs | Syn SNPs | Indels | IG | Mutations per kb | Ns SNPs/indels per kb |
| --- | --- | --- | --- | --- | --- | --- | --- |
| Pyoverdine | 72.2 | 173 | 93 | 38 | 8 | 4.32 | 2.92 |
| Pyochelin | 24.937 | 63 | 34 | 12 | 7 | 4.61 | 3.01 |
| *phu* | 7.516 | 31 | 10 | 3 | 34 | 10.38 | 4.52 |
| *has* | 10.8 (8.8) | 17 | 14 | 2 |  | 3.05 (3.75) | 1.75 (2.16) |
| *feo* | 2.19 | 6 | 3 |  |  | 4.10 | 2.74 |

